# HolistIC: Leveraging Hi-C and Whole Genome Shotgun Sequencing for Double Minute Chromosome Discovery

**DOI:** 10.1101/2021.03.22.435038

**Authors:** Matthew Hayes, Angela Nguyen, Rahib Islam, Caryn Butler, Ethan Tran, Derrick Mullins, Chindo Hicks

## Abstract

Double minute chromosomes are acentric extrachromosomal DNA artifacts that are frequently observed in the cells of numerous cancers. They are highly amplified and contain oncogenes and drug resistance genes, making their presence a challenge for effective cancer treatment. Algorithmic discovery of double minutes (DM) can potentially improve bench-derived therapies for cancer treatment. A hindrance to this task is that DMs evolve, yielding circular chromatin that shares segments from progenitor double minutes. This creates double minutes with overlapping amplicon coordinates. Existing DM discovery algorithms use whole genome shotgun sequencing in isolation, which can potentially incorrectly classify DMs that share overlapping coordinates. In this study, we describe an algorithm called “ HolistIC” that can predict double minutes in tumor genomes by integrating whole genome shotgun sequencing (WGS) and Hi-C sequencing data. The consolidation of these sources of information resolves ambiguity in double minute amplicon prediction that exists in DM prediction with WGS data used in isolation. We implemented and tested our algorithm on the tandem Hi-C and WGS datasets of three cancer datasets and a simulated dataset. Results on the cancer datasets demonstrated HolistIC’s ability to predict DMs from Hi-C and WGS data in tandem. The results on the simulated data showed the HolistIC can accurately distinguish double minutes that have overlapping amplicon coordinates, an advance over methods that predict extrachromosomal amplification using WGS data in isolation.

**Availability:** Our software is available at http://www.github.com/mhayes20/HolistIC.

## 1 Introduction

Double minute chromosomes (DM) are extrachromosomal, circular, acentric fragments of DNA that are found in the cells of various cancer subtypes [4, 27]. Their mechanism of formation is unclear, but hypothesized models suggest that they can form as a result of chromothripsis [21], breakage-fusion-bridge cycles [23] or episomal insertion of DNA [22]. Since DMs lack centromeres, they are randomly passed to daughter cells during cell division, a process that increases their numbers in tumor cells. Accordingly, DMs are highly amplified, often with dozens of copies in tumor cells. Furthermore, they contain oncogenes, giving tumor cells a survival advantage. Thus, the elimination of DMs from cancer cells has important implications for the amelioration of cancer malignancy. Studies have explored approaches to reduce the number of DMs [27], with one study indicating that a certain chemotherapy drug can help to reduce their numbers. Other bench studies have sought to reduce the their numbers by inducing micronuclei formation [11].

Since cancer therapies can be aided by the reduction of DMs in tumor cells, it is important to develop methods to discover their presence in cancer genomes. Discovery of double minutes can be performed algorithmically using whole genome shotgun sequencing data (WGS). Methods that use WGS data for automated DM discovery must perform the following steps: 1) acquire the breakpoints of contiguous genomic regions with amplified copy number, 2) acquire the breakpoints of ostensible structural variants, and 3) using the information gathered from steps 1 and 2, consolidate it to make a final list of predictions for DM amplicons.^5^

Since DMs are highly amplified, their distinct segments should have elevated copy number (i.e. read depth) in the WGS data. Furthermore, the breakpoints between segments will resemble structural variant signals (though these breakpoints may not truly represent such events). One method for addressing this problem, DMFinder, predicts double minutes by incorporating external predictions for amplicons and structural variant breakpoints. It creates a graph where vertices are amplicons and edges are predicted breakpoints between amplicons [10]. After locating all strongly connected components, the double minutes are predicted as the set of cycles in each component. If no cycles are found, the method searches for chains of amplicon vertices connected by edges. The AmpliconFinder algorithm [17] is designed to find double minute amplicons in isolation, but not the breakpoint adjacencies. This algorithm is based on a hidden Markov Model; windows of a fixed size are labeled as 0 or 1, depending on their read depth. Regions of high local read depth (i.e. clusters of 1s) are predicted as amplicons after solving a system of equations related to the model. Another algorithm, AmpliconArchitect, can predict extrachromosomal amplified DNA using WGS data [7]. The method takes as input a set of aligned reads and a set of seed intervals. The method merges seed intervals and identifies amplicon breakpoints by finding positions with sharp changes in copy number. After a breakpoint refinement step, it generates a breakpoint graph used to refine amplicon copy counts. The method decomposes the graph into simple cycles, indicating a putative variant of interest. The method also allows for simple cycles to be merged, allowing for multiple amplicon architectures to be considered. Another algorithm for this problem, called CouGaR, can identify complex rearrangements, including double minutes, using WGS data [8]. Like the other methods, CouGaR uses WGS data to identify amplicons connected by structural variant breakpoints. The method first compiles a list of tumor adjacencies, followed by a prediction of amplified regions. It uses this information to build a tumor adjacency graph, subsequently predicting tumor copy number by formulating the tumor adjacency graph as a network circulation problem. Lastly, final predictions are made by taking the predicted amplified tumor contigs and finding the simple cycles in the graph that can best explain the data. The aforementioned algorithms share a common limitation: they rely only on short read whole genome sequencing data (WGS) to make their predictions. Using WGS data in isolation is limited for accurate discovery of double minute chromosomes. Double minutes can evolve as cancer progresses, yielding derivative DMs with shared amplicons [26]. Since WGS data presents the genome linearly, it is difficult to distinguish a DM with many amplicons from multiple DMs with overlapping amplicons. This is depicted in Figure 1. For example, using only whole genome WGS data, it is difficult to determine if the red-green-blue-yellow-red-green-orange-purple cycle is because of a single “ super” DM (in Figure 1C) or because of the two distinct DMs in Figures 1A and 1B with common amplicons (red and green). Only one of these possibilities is the biological truth. However, the ambiguity makes accurate reconstruction difficult in WGS data.

**Fig. 1.**
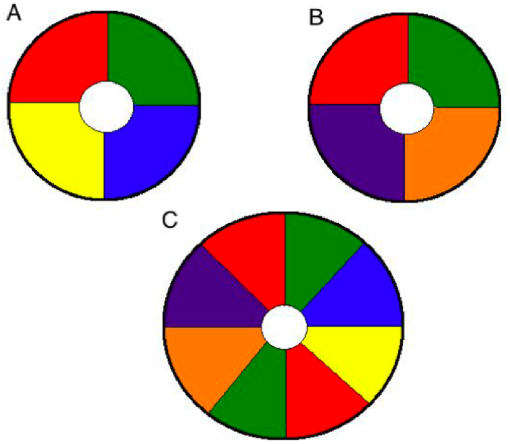
The problem of ambiguity in predicting DMs. The red and green segments in DMs A and B are shared by the DM in panel C.

## 2 Approach

We present an algorithm named HolistIC that can enhance double minute chromosome predictions by predicting DMs with overlapping amplicon coordinates. The method unambiguously resolves DM structure by taking as input a) WGS-predicted double minute amplicon coordinates, b) WGS-predicted breakpoints between amplicons, and c) Hi-C sequencing data from the same sample. HolistIC can uncover double minutes, even in the presence of DM segments with overlapping coordinates. We evaluate our method on the WGS and Hi-C data of three cancer cell lines with DMs: PC3 (prostate cancer), PANC1 (pancreatic cancer), NCIH460 (non-small-cell lung cancer). Figure 2 was generated with JuiceBox [20] and it shows the heat map induced by the Hi-C data of NCIH460. Since double minutes are numerous and randomly scattered throughout a tumor cell, they have elevated genome-wide interaction. Thus, the Hi-C map shows this “ cross” like pattern (green circle) in the NCIH460 cell line, indicating significant interaction with chromatin throughout the genome. We also evaluate the method on simulated datasets consisting of Hi-C reads and whole genome sequencing reads. Figure 3 provides a flowchart describing the approach.

**Fig. 2.**
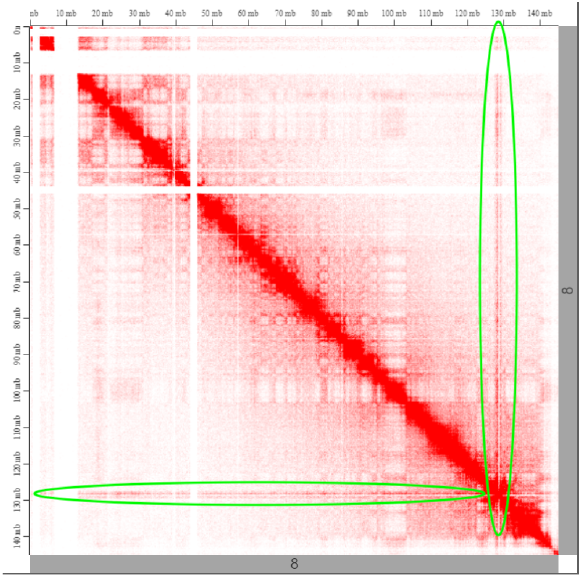
Hi-C contact map for a section of chromosome 8 for the NCIH460 dataset. There is an apparent double minute amplicon at around 130mb (circled in green).

**Fig. 3.**
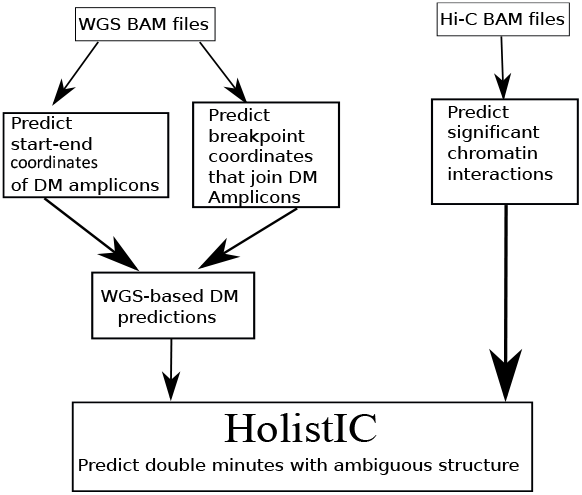
Flowchart of pipeline and how the HolistIC algorithm fits in.

## 3 Methods

### 3.1 Predicting DMs from Whole Genome Sequencing

HolistIC requires as input a list of double minute chromosome predictions, ostensibly from a WGS-based algorithm. We used two methods for this purpose: AmpliconArchitect [7] and DMFinder [10]. DMFinder requires as input a list of predicted contiguous amplified regions. We used AmpliconFinder [17] for this purpose. Furthermore, DMFinder also requires a list of structural variant predictions as input to predict the boundaries between adjacent amplicon intervals. We use Delly [19] for this purpose.

### 3.2 Validating DMs using Hi-C sequencing data

Once we have a set of DMs predicted from WGS data, we can use Hi-C data from the same sample to resolve possible ambiguities in DM predictions from the WGS data. We can do so using Algorithm 1:

### 3.3 Definitions

Algorithm 1 uses the following functions and sets:

#### Algorithm 1 Algorithm to distinguish DMs with shared amplicon coordinates

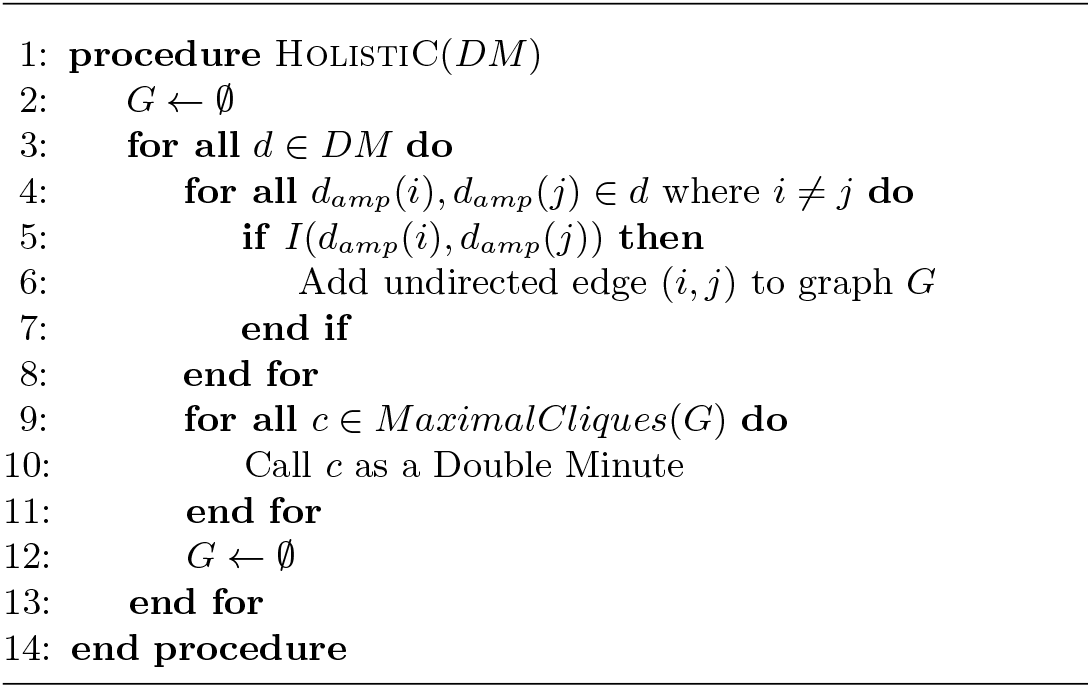

- *DM* : the input set of predicted double minutes from WGS data
- *d*_*amp*_(*i*): refers to amplicon interval *i* that belongs to a double minute *d*.
- *MaximalCliques*(*G*) : After building the graph in steps 4-8, this set contains the maximal cliques of the resulting graph.
- *I*(*x, y*) : A Boolean function that returns **true** if there is significant Hi-C interaction between amplicon intervals *x* and *y*. It returns **false** otherwise.

### 3.4 Explanation of algorithm

Regarding the algorithm, in lines 3-8 it builds a graph *G* where the vertices are predicted double minute amplicon segments (from the WGS data) and an undirected edge between vertices indicates that there is significant Hi-C interaction between the amplicons. We use the GOTHiC and Fit-Hi-C programs to measure significant chromatin interaction in Hi-C data [18, 3]. This allows us to implement the *I*(*x, y*) function required by the algorithm.

In lines 9-10, the algorithm is extracting all maximal cliques identified from the resulting graph *G*. Although maximal clique enumeration is an NP-hard problem, we use the Python library Networkx’s implementation of the Bron-Kerbosch algorithm for this task [2]. This is an exact algorithm to solve the maximal clique problem, but it is fast in practice [5]. We consider each maximal clique a true double minute because we expect there to be significant pairwise chromatin interaction for the amplicons that comprise a DM. This is expected since these amplicons are proximal to one another. Supplementary Tables 1-5 and Supplementary Figures 1 and 3 provide analyses of intra-DM segment interaction from Hi-C cancer data, so this notion is supported by real data. In the case of distinct double minutes with overlapping amplicons, this approach can theoretically allow double minutes to be accurately enumerated since amplicons that are **not** shared among several distinct DMs will not have this heighted Hi-C interaction, and thus will not have edges between their vertices in the graph *G*. Figures 4 and 5 illustrate this. In Figure 4, the double minute induces the interaction graph in this figure. Since all of the amplicons are part of the same DM, they should have heightened pairwise interaction in the Hi-C data. Since all pairs of amplicons have significantly high interaction, the graph should theoretically be complete. After predicting the double minutes from the WGS data, applying the maximal clique algorithm should confirm the true structure of the DM since complete graphs are maximal cliques. In Figure 5, there are two distinct DMs with shared amplicons (green and red). Because of this, using only WGS data, we cannot distinguish between the DMs in Figure 5 and the large DM in Figure 4. For example, assuming the two DMs in Figure 5, when forming the interaction graph during steps 4-6 in the algorithm, the orange and yellow vertices should not be connected by an edge since they are not part of the same DM (i.e. they should have no significant chromatin interaction). Thus, when collecting the maximal cliques from the graph in Figure 5, we should see two maximal cliques, one for the red-blue-green-yellow DM, and one for the red-green-orange-purple DM.

**Fig. 4.**
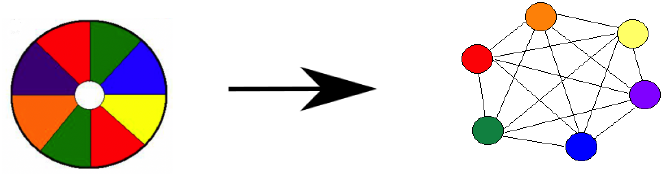
The double minute on the left induces the interaction graph on the right. Since this DM implies a complete graph, the graph is trivially a maximal clique.

**Fig. 5.**
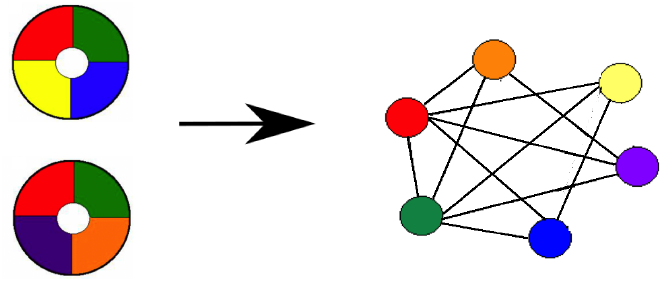
The double minutes on the left induce the interaction graph on the right. Each distinct double minute should induce its own maximal clique.

## 4 Results

To assess our framework, we acquired the WGS and Hi-C sequence data for the PANC1, PC3, and NCIH460 cell lines. These cell lines are known to contain double minute chromosomes [6, 25]. The WGS and Hi-C reads are Illumina paired read datasets with the following accession numbers: PANC1 (SRA accession SRX5466647, ENCODE Project accession #’s ENCFF817XOP, ENCFF876LKL), NCIH460 (SRA accession SRX5466693, ENCODE Project accession #’s ENCFF902YCE, ENCFF665JQY), PC3 (SRA accession SRX5055020, SRX1308509). For the WGS datasets, read duplicates were removed with the MarkDuplicates tool of the Picard Suite [1].

### 4.1 Processing of cancer data

We aligned the aforementioned datasets to the human reference genome hg38. For the WGS data, we aligned it using BWA [16]. For the Hi-C reads, they were processed using the HiCUP program [24] to not only remove common Hi-C sequencing artefacts, but to align the data as well. HiCUP requires use of either Bowtie or Bowtie2 [13, 14] to map reads to a reference; we chose Bowtie2 for these experiments.

Double minute architecture was predicted in the WGS data with the following programs and frameworks:

– AmpliconArchitect [7]
– DMFinder (using AmpliconFinder and Delly results as input) [10, 17, 19].

We used AmpliconArchitect as part of this experiment because it not only allowed us to compare its performance to our WGS-based framework, but it allowed us to gauge the degree to which its predictions were enhanced by the Hi-C capabilities of HolistIC. WGS results from AmpliconArchitect were given as input to the HolistIC algorithm, in addition to Hi-C data results on the same cancer cell lines (described in the next paragraph).

Regarding the DMFinder+AmpliconFinder framework, double minute amplified segments were predicted with AmpliconFinder and amplicon breakpoints were predicted with Delly. The output from Delly and AmpliconFinder was used as input to DMFinder. The DMFinder results were then given as input to HolistIC.

To generate the Hi-C results, we used the programs GOTHiC and Fit-Hi-C [18, 3] to measure significant chromatin interactions in the PC3, NCIH460, and the PANC1 cell lines ^6^. For each WGS-predicted double minute, interactions in these programs were measured for all pairs of Hi-C bins that contained predicted amplicon coordinates. For GOTHiC, bin sizes of 200,000 bp were used, and interactions were counted as significant if the q-value was*≤* 0.05. For Fit-Hi-C, bins were determined by restriction fragments cleaved by BglII (for PC3) and HindIII (for NCIH460 and PANC1). Significant interactions in Fit-Hi-C were those with a q-value*≤* 0.05. After acquiring the GOTHiC and Fit-Hi-C results for all three cell lines, we used these results as input to HolistIC, thus generating the following result sets:

– HolistIC results on AmpliconArchitect (WGS) and GOTHiC (Hi-C)
– HolistIC results on DMFinder+AmpliconFinder (WGS) and GOTHiC
– HolistIC results on AmpliconArchitect (WGS) and Fit-Hi-C (Hi-C)
– HolistIC results on DMFinder+AmpliconFinder (WGS) and Fit-Hi-C (Hi-C)

### 4.2 Simulated data generation

#### Whole genome sequence data

To further evaluate the framework we generated datasets consisting of simulated double minute chromosomes in 1) WGS data and 2) Hi-C data. To generate the WGS data, we created amplicons that included the MYC gene (100 kb upstream and downstream), along with 5 additional randomly-selected genomic intervals, chosen to exclude the centromere, telomere, and regions of heterochromatin. The lengths of these segments were uniformly distributed between 300 kb and 3 Mb, which is reasonable considering the typical length of true biological DM amplicons. After selecting these regions, they were concatenated and amplified 50-fold and 100-fold. For each amplification level, separate FASTA files were created by concatenating the amplified regions to the original reference genome. Four additional DMs were created in this manner, resulting in all of them sharing a segment with the same coordinates (to evaluate the ability of our framework to capture distinct DMs with overlapping amplicon coordinates). In this case, all four DMs shared the MYC amplicon. After creating these modified FASTA files, it was given as input to WGSim [15] with 400-bp fragments, 70-bp standard deviation of fragment lengths, 150-bp paired, reads, and approximately 40X coverage. The data was then aligned to the human reference genome hg19 with BWA. We aligned to hg19 to promote compatibility with AmpliconFinder. DM segment breakpoints were predicted with Delly, while amplicon segments were predicted with AmpliconFinder. The output from these programs was used as input to AmpliconArchitect and to the DMFinder+AmpliconFinder framework. The details are provided in Section 4.2.3.

#### Hi-C sequence data

We used the AveSim program to create simulated Hi-C sequencing reads from the aforementioned 50-fold simulated WGS data [6] ^7^. This program is used to create simulated Hi-C data that contains an assortment of genomic variant types, including double minutes. However, AveSim only simulates chromatin interactions for genomes that contain a single double minute with a single amplicon segment. This is not sufficient for our needs since our algorithms assume that genomes can contain multiple DMs, each with multiple segments (which is also plausible since this is seen in many cancer genomes). Following a recommendation from the developer of AveSim, we altered the AveSim framework as follows: we first used the modified FASTA file from the 50-fold WGS experiment and created a custom BSgenome object from the Bioconductor package. This object was used as the default genome for AveSim since it contained the simulated DMs of interest. For each double minute amplicon segment generated in the WGS step, AveSim was executed separately. The resulting FASTQ files were concatenated to form the complete set of simulated Hi-C reads. These reads were processed and aligned to the the human reference genome hg38 with HiCUP. Like the cancer data, GOTHiC and Fit-Hi-C were used to measure chromatin interaction in the Hi-C data. For GOTHiC, We used bin sizes of 200,000 bp and counted pairwise bin interaction as significant if the corresponding q-value was *≤*0.05. For Fit-Hi-C, we used non-fixed bin sizes where the fragments were induced by *Hin*dIII restriction sites.

#### Evaluation

The aforementioned datasets were provided as input to HolistIC, pursuant to the following list:

– HolistIC results on AmpliconArchitect (WGS) and GOTHiC (Hi-C)
– HolistIC results on DMFinder+AmpliconFinder (WGS) and GOTHiC
– HolistIC results on AmpliconArchitect (WGS) and Fit-Hi-C (Hi-C)
– HolistIC results on DMFinder+AmpliconFinder (WGS) and Fit-Hi-C (Hi-C)

For each WGS dataset, results on the 50-fold and 100-fold results were reported separately to assess the ability of each approach to reconstruct DM amplicons in varying amplification scenarios.

### 4.3 Results on Simulated Data

In our first experiment, we sought to determine if AmpliconArchitect [7] and DMFinder could accurately predict the correct structure of the simulated double minutes, given that they share a common segment on chromosome 8. We expected that this ambiguity is difficult for WGS-only methods to resolve. Our results confirmed this, as neither method correctly resolved this structure when used in isolation, as shown in Figure 6, Table 1, and Supplementary Table 7. In these tables, each row is a DM amplicon. Each simulated double minute is identified in the leftmost column, along with their amplicon coordinates in columns 2-4. For all of the blue “ N” entries in columns 5 and 8, DMFinder and AmpliconArchitect predicted all of these amplicons as belonging to the same double minute; this is erroneous, but is expected since they only use whole genome sequencing data. Also in these columns, the amplicon intervals are labeled Y if the WGS-only method successfully associated them with their correct DM. Columns 6-7 and 9-10 provide the ID of the maximal clique identified by HolistIC that contains the amplicon.

**Table 1.**
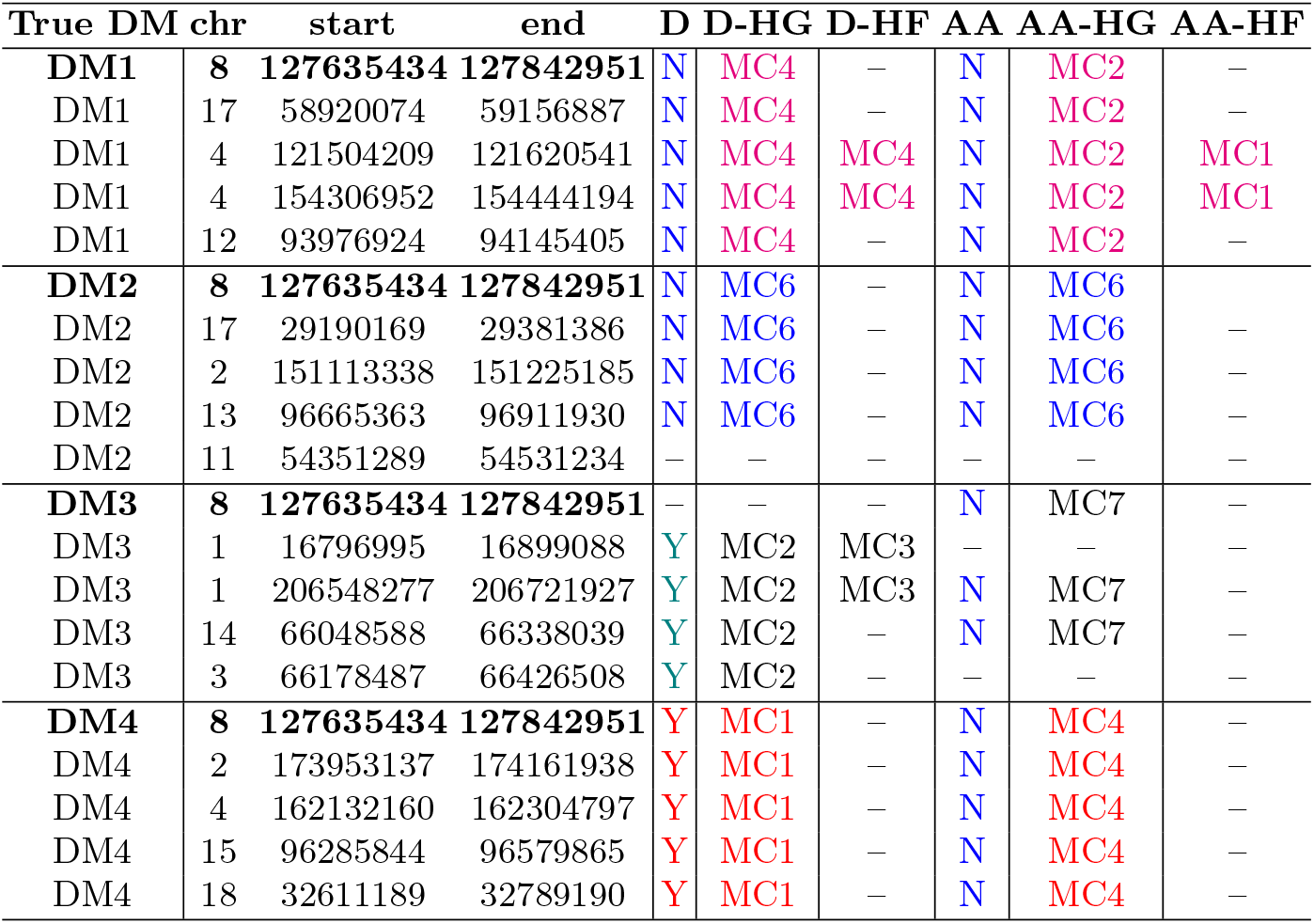
Results on the 50-fold amplification dataset. The table shows the simulated double minutes, their amplicon interval coordinates, and the results of each method for each interval. Legend: D - DMFinder, D-HG – HolistIC (DMFinder+GOTHiC), D-HF - HolistIC (DMFinder+Fit-Hi-C), AA - AmpliconArchitect, AA-HG - HolistIC (Am-pliconArchitect+GOTHiC), AA-HF - HolistIC (AmpliconArchitect+Fit-Hi-C). Columns 5 and 8 provide the results of the WGS-only methods DMFinder (D) and AmpliconArchitect (AA); they say whether or not each method correctly associated the amplicon with its true DM. Columns 1,2,4, and 5 indicate the maximal cliques (MC) identified by by each method where the intervals are found. In columns 5-10, uniform colors indicate the intervals that belong to the same double minute (one for each color). Dashes in these columns mean that the method did not predict the amplicon at all. The bolded text in columns 1-4 indicate coordinates of the MYC amplicon that is shared among all DMs.

**Fig. 6.**
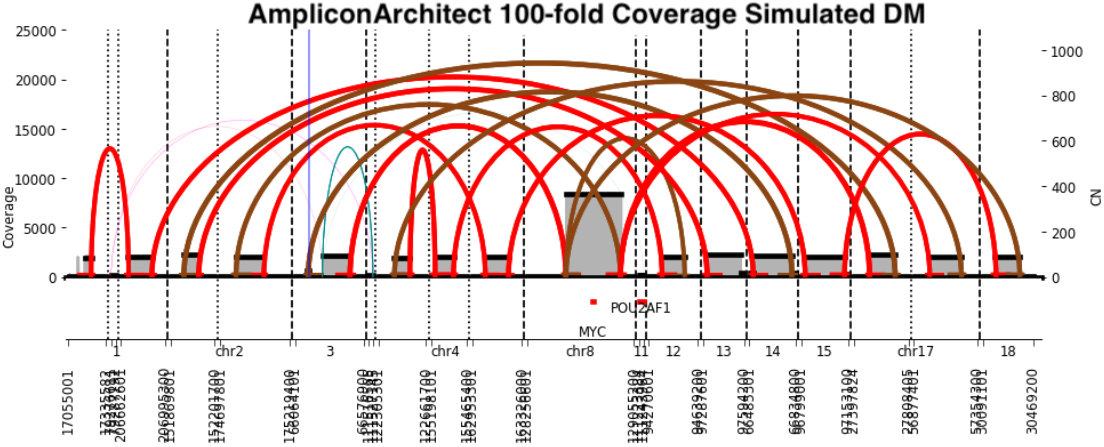
AmpliconArchitect prediction for the four double minutes in the 100-fold simulated dataset. The shared MYC amplicon is evident. AmpliconArchitect erroneously predicted this as a single “ super” double minute. The image was produced by AmpliconArchitect.

AmpliconArchitect predicted intervals in all of the double minutes as belonging to a single super DM (Figure 6 shows a prediction for the 100-fold simulated data). DMFinder partially segregated the intervals into their correct DM, but not for the true DMs MC1 and MC2. This lack of precision is ostensibly a consequence of WGS-only prediction. To address this, we used these WGS intervals as input to HolistIC, combined with the Hi-C data for this simulated genome. Significant Hi-C interactions were measured with GOTHiC and Fit-Hi-C; the results were gathered for each Hi-C method separately. When using GOTHiC, HolistIC accurately segregated the double minute intervals into their correct structure for both AmpliconArchitect and DMFinder, with a few exceptions. This was evident for both the 50-fold and 100-fold amplification datasets.

Specifically for DMFinder, we used GOTHiC and Fit-Hi-C as input to HolistIC (in separate columns in Table 1 and Supplementary Table 7). In these experiments, we measured the number of multiple predicted amplicons, incorrectly assigned to a “ super” double minute by DMFinder alone, that were correctly predicted as smaller, individual double minutes by HolistIC. Figure 7 provides the HolistIC interaction graph for one of the simulated double minutes identified by DMFinder. Each vertex is an amplicon identified by AmpliconFinder ^8^; the set of vertices were confirmed as a double minute by HolistIC since the graph is complete and thus induces its own maximal clique. Since the original double minute consisted of this exact set of vertices, there is complete concordance between the HolistIC results and the DMFinder results on this double minute, despite the fact that the chromosome 8 amplicon is shared by three other simulated DMs in the dataset. However, Supplementary Figure 2 depicts the interaction graph constructed by HolistIC for just **one** of the predicted double minutes by DMFinder. According to the interaction graph, there is not significant interaction between all pairs of vertices, which is **not** indicative of a single double minute. In fact, this graph depicts four separate maximal cliques, all of which share the chromosome 8 simulated amplicon in the middle. Although DMFinder predicted these vertices as all belonging to the same double minute, this graph is actually comprised of **three** of the simulated double minutes. In this same graph, the true positive results of HolistIC are overlaid. Each of the maximal cliques identified by HolistIC are circled in this graph, which are the program’s predictions for the subgraphs that denote each DM. The pink subgraph represents a DM that was perfectly identified since all five original amplicons were present. Four of the amplicons were identified for the DM’s subgraph circled in blue, while just two amplicons for the green subgraph were identified.

**Fig. 7.**
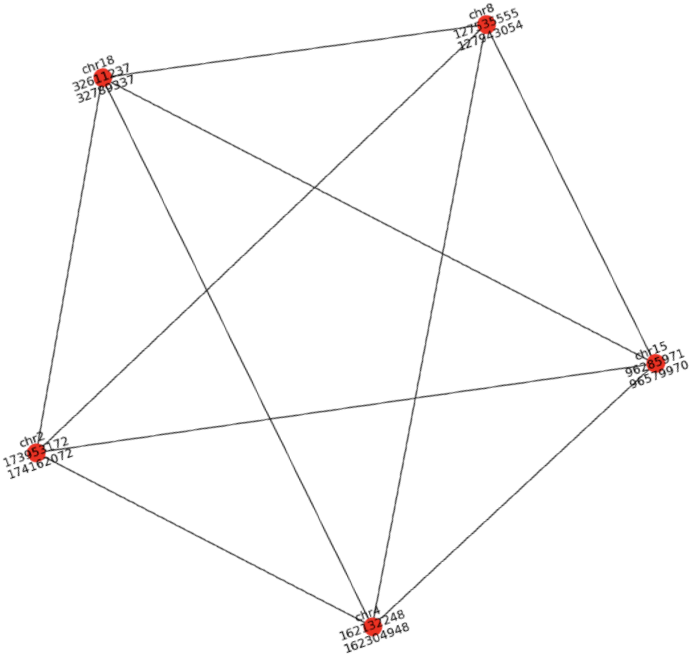
The interaction graph constructed by HolistIC for one of the true simulated double minutes. All five amplicons are presented and all pairs of amplicons show heigtened interaction in the Hi-C data, which is expected for double minutes.

For AmpliconArchitect, there were similar results for the simulated data, for both the 50-fold and 100-fold datasets. AmpliconArchitect was first applied to the simulated data and the results were gathered without applying them to HolistIC. Although AmpliconArchitect identified nearly all of the true amplicon intervals belonging to the simulated double minutes, it erroneously predicted them as belonging to a single super DM, as shown in Table 1 and Supplementary Table 7. However, after applying HolistIC by using GOTHiC results, there is clearer separation of the intervals into their correct DM.

HolistIC lost sensitivity when using the Fit-Hi-C data for both DMFinder and AmpliconArchitect. This is likely because Fit-Hi-C is optimized for medium to long range chromatin interactions which could cause shorter range interactions within DM amplicons to be missed. As a result, there are a large number of false negatives when using Fit-Hi-C for the HolistIC framework.

There were several false positive maximal cliques in these results. The clique identified as “ MC3” is an example. This clique was identified by HolistIC+DMFinder+GOTHiC (D-HG) but is spurious since it is comprised of amplicons that do not all belong to the same true DM. MC3, shown in Supplementary Table 6, was predicted because of the *cis*-interaction between the chromosome 17 segments; intrachromosomal interactions were elevated in the simulated data which caused an interaction edge to be placed between these segments, thus inducing a spurious maximal clique. There is another redundant clique (MC5) not listed in Supplementary Table 6. However, the MC5 clique actually contains two amplicons that are part of a true double minute MC2. This redundant prediction was due to the imprecise prediction by the DMFinder framework of the chromosome 3 interval. The AA-HG framework presented a similar number of false positives on the simulated data due to the same reasons. This was true for the 50-fold and 100-fold datasets.

#### Interval segregation without adjacency information

The extraction of information required to predict extrachromosomal circular DNA is difficult to perform and relies on accurate prediction and classification of amplicon coordinate boundaries. In our experiments, we used Delly to predict amplicon adjacencies. However, in case this information is not available, we assessed the ability of HolistIC to accurately segregate predicted amplicons into their proper double minute. For this experiment, we used the DMFinder results from Table 1, except that we falsely changed the labels of all predicted amplicons to belong to a single double minute: DM1. This has the effect of removing the amplicon adjacency information which is needed to build the double minute using only WGS data. As a result of this procedure, each predicted amplicon is initially assumed by HolistIC to belong to a single DM. Although this is not the reality, it was performed to assess HolistIC’s ability to accurately segregate the amplicons by their true DM using **only** the set of predicted amplicons and **not** prior information about the amplicon’s assigned DM. Figure 8 provides the interaction graph constructed by HolistIC from these data, with the maximal cliques identified by the colored circles. From this graph, it is evident that there are at least four maximal cliques that represent double minutes with a common MYC amplicon at chromosome 8. Despite not having the inter-amplicon breakpoint information, HolistIC still accurately separated amplicons into their proper cliques, and thus their proper double minutes. There was a false positive maximal clique due to the edge that connects the chromosome 17 vertices in the purple and blue components. In general, intrachromosomal, or *cis-*interactions are more common in Hi-C data due to chromosome territories [12]. This phenomenon could increase the likelihood of spurious contacts between intrachromosomal loci, which could promote false positives.

**Fig. 8.**
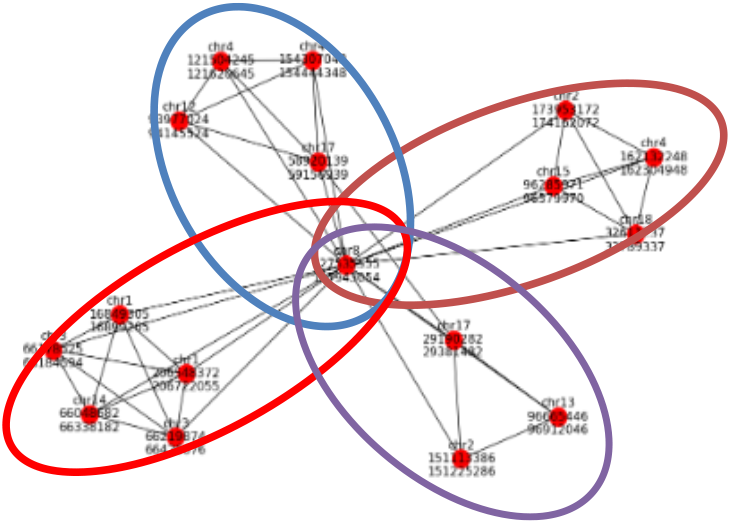
The interaction graph visualized by HolistIC for the amplicons listed in Table 1. Despite initially being labeled as belonging to the same DM, HolistIC accurately separated the amplicons into their true DMs (circled).

### 4.4 Results on Cancer Data

We applied the evaluation framework to the PC3, PANC1, and NCIH460 datasets; we appled DMFinder and AmpliconArchitect to the cancer data and then applied HolistIC to the results of these methods. For PANC1, there is a double minute chromosome with an amplicon interval at chr19:39,774,739–40,847,406 (in hg19 coordinates). For NCIH460 there is a double minute chromosome with an amplicon interval at chr8:128,691,158–130,402,069. For the DMFinder framework (D-HG) on PANC1 and NCIH460, AmpliconFinder predicted a large number of isolated amplified segments in each dataset (71 in PANC1, 85 in NCIH460; Table 4). However, DMFinder subsequently failed to predict the amplified intervals as circular or linear amplified contigs. This may be due to insufficient simple structural variant signals between amplicon breakpoints (which would cause “ isolated” amplified regions not connected by SV breakpoints). Consequently, if the amplicon breakpoints are not accurately predicted, then DMFinder can also lose sensitivity. Thus, for PANC1 and NCIH460, we used AmpliconFinder+HolistIC+GOTHiC (AF-HG) instead of DMFinder. Since AmpliconFinder only finds amplified intervals and not their adjacencies, we treated the predicted intervals as belonging to a single double minute. This assessed the ability of HolistIC to separate “ ‘isolated” amplicons into their true DMs. Fit-Hi-C failed to yield significant chromatin contacts for the cancer data so it was omitted from our analyses. The results are presented in Tables 2 and 3.

**Table 2.**
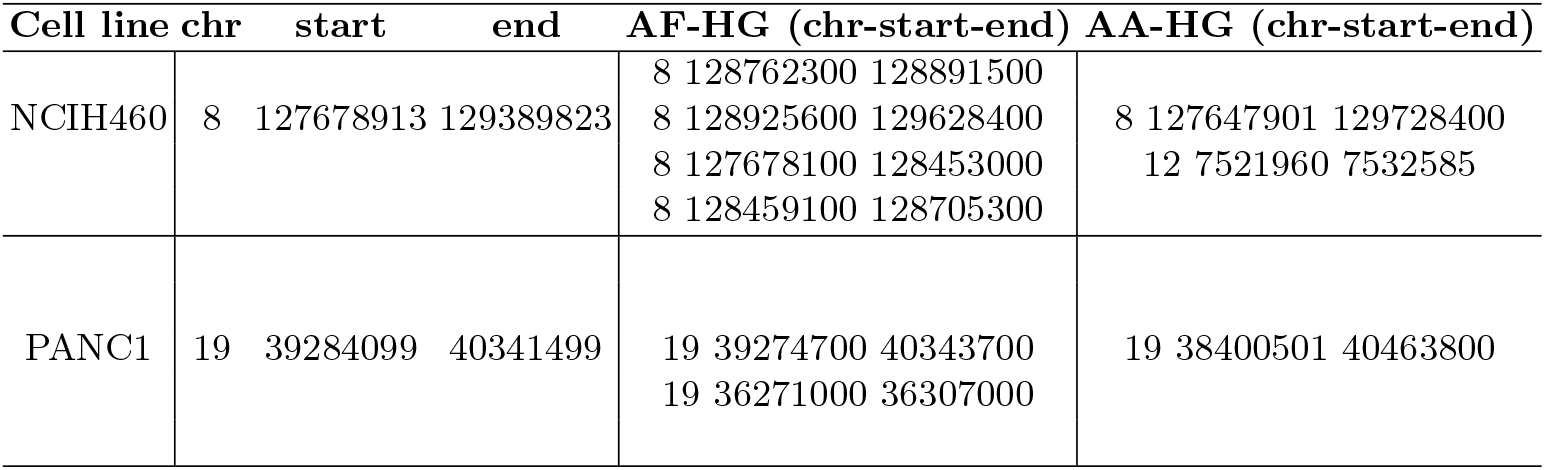
Results on the cancer cell line datasets (hg38 coordinates). Columns 2-4 contain the ground truth interval coordinates. The PANC1 amplicon was the only interval predicted by AmpliconArchitect so it is trivially its own maximal clique.

| Cell line | chr | start | end | AF-HG (chr-start-end) | AA-HG (chr-start-end) |
| --- | --- | --- | --- | --- | --- |
| NCIH460 | 8 | 127678913 | 129389823 | 8 128762300 128891500 | 8 127647901 129728400<br>12 7521960 7532585 |
|  |  |  |  | 8 128925600 129628400 |  |
|  |  |  |  | 8 127678100 128453000 |  |
|  |  |  |  | 8 128459100 128705300 |  |
| PANC1 | 19 | 39284099 | 40341499 | 19 39274700 40343700 | 19 38400501 40463800 |
|  |  |  |  | 19 36271000 36307000 |  |

**Table 3.** Results on the PC3 cell line for the AF-HG pipeline. Columns 2-4 contain the ground truth interval coordinates. The colored rows are distinct predicted DMs. The red DM is a potential false positive.

| Cell line | chr | start | end | AF-HG (chr-start-end) |
| --- | --- | --- | --- | --- |
| PC3 | 8 | 121853389 | 121954300 | 8 133661600 133824200 |
|  | 8 | 132851686 | 132892344 | 8 132851800 132892500 |
|  | 8 | 127717002 | 127818977 | 8 126554700 126641500 |
|  | 8 | 133661572 | 133824065 | 8 126248900 126264100 |
|  | 8 | 130455524 | 130696951 | 8 137239000 137274000 |
|  | 8 | 141195337 | 141237973 | 8 130455600 130861100 |

**Table 4.** Summary of double minute prediction on cancer data: AmpliconFinder (AF) vs. HolistIC (AF-HG framework).

| Cell line | AF #intervals predicted | AF-HG max. cliques (#vertices) | True amplicon found (AmpliconFinder/HolistIC)? |
| --- | --- | --- | --- |
| PANC1 | 71 | 15 (32) | Y/Y |
| NCIH460 | 85 | 9 (23) | Y/Y |

AmpliconArchitect predicted the amplicon coordinates at high accuracy. The single amplicon interval for PANC1 (Table 2) is trivially its own maximal clique; no other amplicons were predicted by AmpliconArchitect in this dataset. For NCIH460, AmpliconArchitect predicted the ground truth amplicon at chromosome 8, but also an amplified interval at chromosome 12 that it predicted as being part of the same contig. This suggests that the chr8 and chr12 segments are part of the same double minute. HolistIC, under the AA-HG pipeline, confirmed heightened chromatin interaction between these segments, as they were predicted as being part of the same maximal clique (Table 2). Supplementary Table 1 and Supplementary Figure 1 show a larger-than-normal number of chromatin touches between these segments, thus presenting evidence of intra-DM chromatin interaction. The AmpliconFinder pipeline, AF-HG, yielded more false positives than the AA-HG pipeline. This is likely because the AF-HG pipeline only used predicted isolated intervals as input instead of the circular or linear contigs, which would have been predicted by DMFinder. Nonetheless, the AF-HG pipeline still managed to predict maximal cliques that overlapped the true amplicons, despite some spurious predictions in PANC1 (at chr19:36,271,000) and redundant chr8 predictions in NCIH460.

The results on PC3 are in Table 3 (AF-HG pipeline) and Supplementary Table 8 (AA-HG pipeline). The article by Wu et al. [25] used AmpliconArchitect to predict the structure of a double minute in this cell line. These predicted intervals are in columns 2-4. Columns 5-7 present the AmpliconFinder+HolistIC+GOTHiC (AF-HG) pipeline results where each color indicates a separate predicted double minute. The red intervals are a likely false positive since these intervals were not among those from the original study. Furthermore, the blue and pink intervals are classified as belonging to separate double minutes, despite overlapping with the ground truth intervals. Furthermore, after applying HolistIC to the AmpliconArchitect results, a total of 13 double minutes across 41 intervals were predicted. Further, according to Wu et al., the given double minute was one of several candidate architectures. Thus, while the true DM appears to be among the intervals in columns 2-4, it may also include other intervals, or there could be several distinct double minutes. Nonetheless, the number of predicted DMs by the AA-HG framework suggests that Hi-C interaction can help to further refine extrachromosomal circular DNA (eccDNA) architecture predictions by associating proximal amplicon intervals.

## 5 Discussion

The results on PANC1 and NCIH460 show that HolistIC can associate amplified intervals with their true contig when Hi-C data is present. Supplementary Figures 1 and 3 and Supplementary Tables 1-5 present evidence of intra-DM chromatin interaction, suggesting the viability of this feature for eccDNA prediction.^9^. For the PC3 dataset, the double minute was more complex. The AF-HG pipeline predicted 6 total intervals, with 4 overlapping the intervals from Wu et al. The AA-HG pipeline understandably had more overlap, but it predicted a total of 13 double minutes. Further analysis of PC3, including wet lab analysis, may be necessary to more accurately determine the true structure and number of double minute chromosome populations in this cell line.

The results on the simulated data illustrate HolistIC’s ability to accurately cluster amplicons into separate double minutes. For the DMFinder and AmpliconFinder HolistIC pipelines using GOTHiC, the program accurately distinguished the DMs from each other, despite the fact that the they shared an amplicon with common coordinates (which is a problem for WGS-only algorithms, as illustrated by DMFinder and AmpliconFinder used in isolation). The results also show that HolistIC can accurately cluster amplicons into their true DMs, despite there being no information about amplicon breakpoint adjacencies. Although in such cases there would be no information about the orientation or specific structure of the DM, clustering the amplicons to their correct DM may still hold clinical relevance.

One potential drawback of HolistIC is its performance on double minutes containing amplicons from a single chromosome. The Hi-C data for “ cis” interactions may show higher-than-normal interaction for such amplicons, although this was mostly observed in the simulated data. Moreover, HolistIC may lose sensitivity as the double minutes increase in size, due to the decay in contact probability for distal loci [3]. Despite these limitations, HolistIC is ideal for confirming the true association of amplicons to circular extrachromosomal DNA. Furthermore, it is modular in that the double minute prediction input (including the breakpoint discovery and amplicon discovery stages) can be from any program, as well as the Hi-C interaction input. This lends additional flexibility for use as post-processing program for current and future eccDNA discovery algorithms that use short read sequencing data.

## 6 Conclusion

We have presented an algorithm called HolistIC that can predict double minute chromosomes, given predictions for double minute chromosome amplicons and predictions of significant chromatin interaction from Hi-C data. For future work, we will address the issue of lost specificity for “ cis” interactions, although this phenomenon was mostly observed in the simulated data. Future work will also oversee the development of “ built-in” algorithms to predict breakpoints between adjacent amplicons, in case this data is not present at the start of the pipeline. Lastly, another way to increase specificity is to quantify the strength of the double minute touch pattern seen in Figure 2 (i.e. the “ cross-like” pattern). If this is performed for all predicted amplicons, it can help to further filter out spurious calls since falsely-predicted amplicons should not show this pattern in the Hi-C data (and would thus have a lower score for this quantity).

## 8 Acknowledgements

We would like to thank Dr. Abhijit Charkraborty of the La Jolla Institute for Immunology for very helpful discussions and insight.

## Footnotes

5 Steps 1 and 2 are interchangeable

6 Fit-Hi-C is suited for medium-range interactions

7 We only generated simulated Hi-C data from the 50-fold dataset.

8 AmpliconFinder is used as input to DMFinder. It is a program that finds contiguous regions of elevated copy number.

9 Although the Hi-C sequence data of GBM176 and GBM180 (Gene Expression Omnibus accession # GSE81879, [9]) were analyzed in these tables, we did not apply them to the HolistIC framework due to a lack of WGS data

