## Supplementary Figures and Tables for "HolistIC: Leveraging Hi-C and Whole Genome Shotgun Sequencing for Double Minute Chromosome Discovery"

Supplementary Tables and Figures

Table 1: Summary of interactions for the NCIH460 double minute, measured as interactions per million nucleotides. The DM-DM column is the number of Hi-C contacts within the chr8:127678913-129389823 double minute interval (hg38 coordinates). DM-chr counts the number of interactions between the double minute interval and the rest of the genome. The chr-chr column counts all other (non-DM) interactions throughout the chromosome. The DM-DM interactions are substantially greater than the DM-chr interactions.

| NCIH460 | **Interaction class** | DM-DM | DM-chr | chr-chr |
|---|---|---|---|---|
| | **I/1000000 nt** | 67,703 | 165 | 24,433 |

Table 2: Summary of interactions for the PANC1 double minute.

| PANC1 | **Interaction class** | DM-DM | DM-chr | chr-chr |
|---|---|---|---|---|
| | **I/1000000 nt** | 27,225 | 52 | 26,286 |

Table 3: Summary of interactions for the GB176 double minute described in [2]. The estimated double minute interval is chr7:54,750,000-55,750,000 (hg19) which corresponds to an amplification of the *EGFR* gene. According to Harewood et al., this gene is amplified by a double minute in this cell line [2].

| GB176 | **Interaction class** | DM-DM | DM-chr | chr-chr |
|---|---|---|---|---|
| | **I/1000000 nt** | 324,240 | 45 | 4,521 |

Table 4: Summary of interactions for the GB180 double minute described in [2]. The estimated double minute interval is chr7:54,750,000-55,750,000 (hg19) which corresponds to an amplification of the *EGFR* gene. According to Harewood et al., this gene is amplified by a double minute in this cell line [2].

| GB180 | **Interaction class** | DM-DM | DM-chr | chr-chr |
|---|---|---|---|---|
| | **I/1000000 nt** | 60,412 | 45 | 3,042 |

Table 5: Summary of interactions for the GB180 double minute described in [2]. This table corresponds to the amplified chromosome 12 segment described in this study, which resembles a double minute signature. The interval is chr12:58,000,000-72,000,000 (hg19) which roughly corresponds to amplifications of *MYC* and *CDK4* in this cell line.

| GB180 | **Interaction class** | DM-DM | DM-chr | chr-chr |
|---|---|---|---|---|
| | **I/1000000 nt** | 11,819 | 68 | 2,997 |

Table 6: False positive double minute identified by HolistIC. For the D-HG pipeline.

| **chr** | **start** | **end** | **HolistIC Max Clique (MC)** |
|---|---|---|---|
| 8 | 127635434 | 127842951 | MC3 |
| 17 | 58920074 | 59156887 | MC3 |
| 17 | 29190282 | 29381482 | MC3 |

Table 7: Results on the 100-fold amplification dataset.

| True DM | chr | start | end | D | D-HG | D-HF | AA | AA-HG | AA-HF |
|---|---|---|---|---|---|---|---|---|---|
| **DM1** | **8** | **127635434** | **127842951** | N | MC4 | – | N | MC2 | – |
| DM1 | 17 | 58920074 | 59156887 | N | MC4 | – | N | MC2 | – |
| DM1 | 4 | 121504209 | 121620541 | N | MC4 | MC4 | N | MC2 | MC1 |
| DM1 | 4 | 154306952 | 154444194 | N | MC4 | MC4 | N | MC2 | MC1 |
| DM1 | 12 | 93976924 | 94145405 | N | MC4 | – | N | MC2 | – |
| **DM2** | **8** | **127635434** | **127842951** | N | MC6 | – | N | MC6 | |
| DM2 | 17 | 29190169 | 29381386 | N | MC6 | – | N | MC6 | – |
| DM2 | 2 | 151113338 | 151225185 | N | MC6 | – | N | MC6 | – |
| DM2 | 13 | 96665363 | 96911930 | N | MC6 | – | N | MC6 | – |
| DM2 | 11 | 54351289 | 54531234 | – | – | – | – | – | – |
| **DM3** | **8** | **127635434** | **127842951** | – | – | – | N | MC7 | – |
| DM3 | 1 | 16796995 | 16899088 | Y | MC2 | MC3 | – | – | – |
| DM3 | 1 | 206548277 | 206721927 | Y | MC2 | MC3 | N | MC7 | – |
| DM3 | 14 | 66048588 | 66338039 | Y | MC2 | – | N | MC7 | – |
| DM3 | 3 | 66178487 | 66426508 | Y | MC2 | – | – | – | – |
| **DM4** | **8** | **127635434** | **127842951** | Y | MC1 | – | N | MC4 | – |
| DM4 | 2 | 173953137 | 174161938 | Y | MC1 | – | N | MC4 | – |
| DM4 | 4 | 162132160 | 162304797 | Y | MC1 | – | N | MC4 | – |
| DM4 | 15 | 96285844 | 96579865 | Y | MC1 | – | N | MC4 | – |
| DM4 | 18 | 32611189 | 32789190 | Y | MC1 | – | N | MC4 | – |

Table 8: Results on the PC3 cell line for the AA-HG pipeline. Columns 2-4 contain the ground truth interval coordinates. Predicted HolistIC double minutes are in distinct colors. Only those predicted double minutes are listed that overlap with the coordinates in columns 2-4.

| Cell line | chr | start | end | AF-HG (chr-start-end) |
|---|---|---|---|---|
| PC3 | 8 | 121853389 | 121954300 | 8 130335001 130973000 |
| | 8 | 132851686 | 132892344 | 8 132821986 133002344 |
| | 8 | 127717002 | 127818977 | 8 133561001 133934065 |
| | 8 | 133661572 | 133824065 | 8 133561001 133934065 |
| | 8 | 130455524 | 130696951 | 8 141081728 141382086 |
| | 8 | 141195337 | 141237973 | 8 132821986 133002344 |
| | | | | 8 133561001 133934065 |
| | | | | 8 127597001 127883000 |
| | | | | 8 130335001 130973000 |

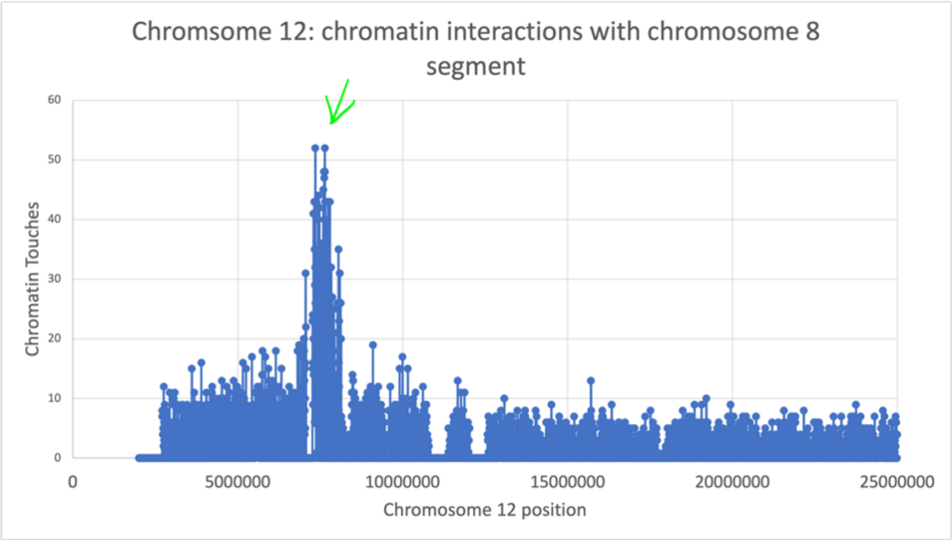


Figure 1: Number of chromatin touches between the NCIH460 chromosome 8 double minute segment, described in [1], and chromosome 12 of this cell line. There is a spike in chromatin touches at approximately 7.5Mb, suggesting that the chromosome 12 amplicon predicted by AmpliconArchitect has heightened interaction with the chromosome 8 double minute amplicon. These segments were predicted by AmpliconArchitect as belonging to the same amplified contig; this was also confirmed by HolistIC. Thus, this figure provides evidence for heightened chromatin interaction within double minute segments. Only the first part of chromosome 12 is shown, but the remaining chromosome 12 loci do not show interaction spikes like that seen in this figure. The loci are grouped into bins of 5,000 nt and the touches were counted directly from the SAM file corresponding to this data.

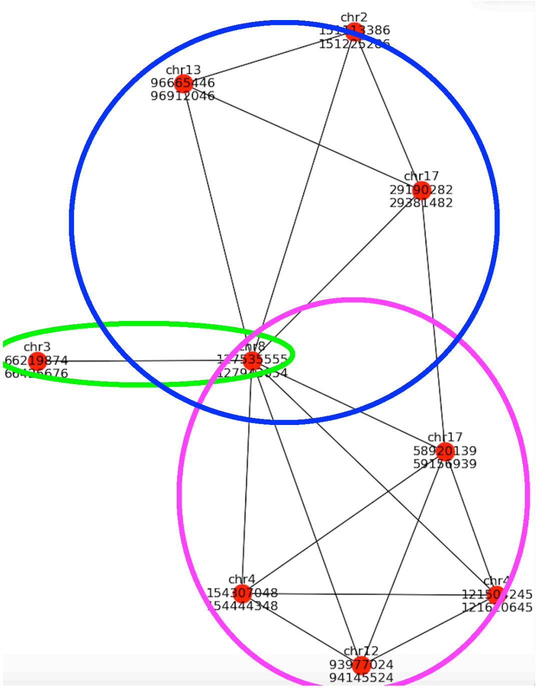


Figure 2: The graph from Figure 8 in the main article, but with maximal cliques circled that were identified by HolistIC. Each of the circled subgraphs indicates a true DM created in the simulated data.

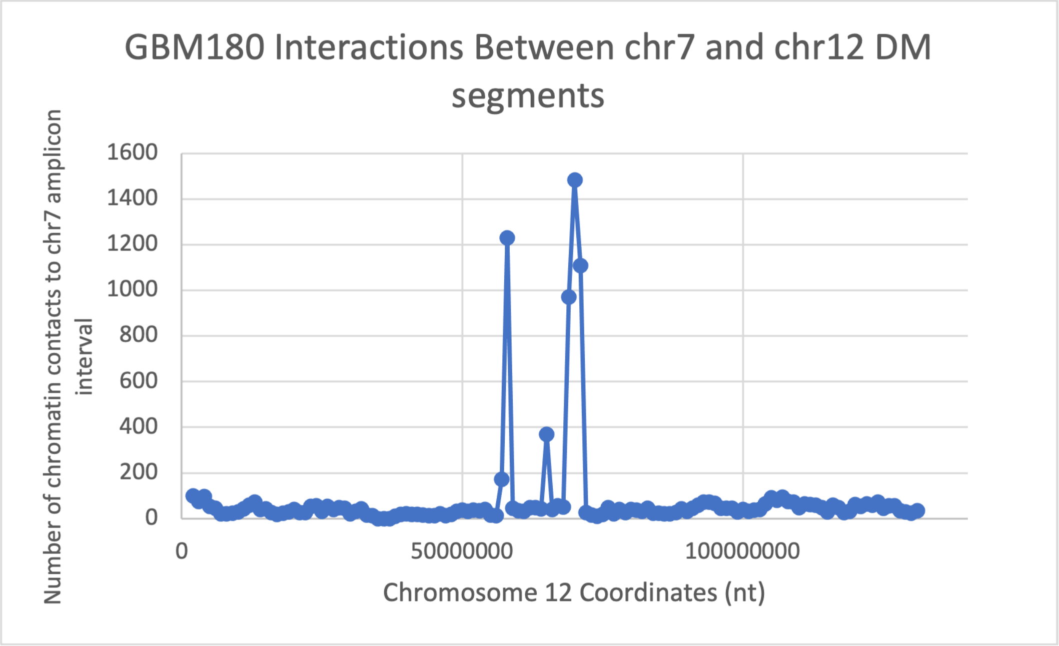


Figure 3: Number of chromatin touches between the GBM180 chromosome 8 double minute segment, described in [2], and chromosome 12 of this cell line. There are spikes in chromatin touches at the chromosome 12 locations at roughly 58Mb and 69Mb. These locations correspond to amplifications of *CDK4* and *MDM2*, respectively. Like Supplementary Figure 1, this suggests intra-DM chromatin interaction, a feature that the HolistIC algorithm uses to make predictions. The loci are grouped into bins of 1,000,000 nt and the interactions were counted from the GB180 interaction data (Gene Expression Omnibus: accession number GSE81879).
